# Three-dimensional super-resolution fluorescence imaging of DNA

**DOI:** 10.1101/796151

**Authors:** Sevim Yardimci, Daniel R. Burnham, Samantha Y. A. Terry, Hasan Yardimci

## Abstract

Recent advances in fluorescence super-resolution microscopy are providing important insights into details of cellular structures. To acquire three dimensional (3D) super-resolution images of DNA, we combined binding activated localization microscopy (BALM) using fluorescent double-stranded DNA intercalators and optical astigmatism. We quantitatively establish the advantage of mono-over bis-intercalators before demonstrating the approach by visualizing single DNA molecules stretched between microspheres at various heights. Finally, the approach is applied to the more complex environment of intact and damaged metaphase chromosomes, unravelling their structural features.

---

Folding of DNA into chromatin is essential for packaging in the nucleus and plays a key role in the regulation of protein-nucleic acid interactions. It is essential to understand chromosome architecture because genome organization has significant impact on cellular processes such as DNA replication, recombination, repair, gene regulation and cell division. Chromosomes are dynamic entities with morphology alterations throughout the cell cycle. During mitosis, human chromosomes adopt a compact X shape before segregation of sister chromatids into daughter cells. Defects in DNA replication, recombination and repair can lead to aberrant chromosome structures manifested by breaks or gaps seen in metaphase spreads^1^. The most commonly employed techniques for investigating chromosome morphology are bright-field and wide-field fluorescence microscopy in which DNA is labelled with a DNA binding probe such as Giemsa or DAPI. While most light microscopy techniques provide two-dimensional (2D) pictures of chromosomes, electron microscopy (EM) and atomic force microscopy (AFM) have been the major methods for the investigation of three-dimensional chromosome structure^2,3^. However, unlike light microscopy in which DNA and DNA-binding factors can be labelled specifically, EM and AFM probe the whole structure of an assembly and cannot differentiate between distinct parts of complex molecules.

Super-resolution fluorescence microscopy methods have become powerful tools for high-resolution structural investigations^4^. An elegant technique to achieve super-resolution imaging of DNA is <u>b</u>inding <u>a</u>ctivated localization <u>m</u>icroscopy (BALM)^5^. BALM relies on binding and dissociation/photobleaching of fluorescent DNA intercalating dyes and the localization from the associated increase in signal with high resolution. While the fluorophore signal intensity is important for localization precision, higher DNA association and dissociation rates are desired to increase the localization density in super-resolution images per unit time. A variety of DNA intercalators and buffers have been tested previously and YOYO-1, a double stranded DNA (dsDNA) intercalating dye, in combination with ROXS (ascorbic acid and methyl viologen) - containing buffer was used for optimal imaging conditions^5^. We compared binding and dissociation kinetics of YOYO-1 and SYTOX Orange (SxO), another dsDNA intercalator commonly used in single-molecule studies^6-8^, under different buffer conditions. Using an autocatalytic model for association of both dyes demonstrates that SxO associates faster due to higher autocatalysis, that is, DNA bound SxO acts to cooperatively bind additional dye at a greater rate than YOYO-1 (Fig. 1a,b and Supplementary Fig. 1a). YOYO-1 and SxO can be modelled as dissociating with mono and bi-phasic kinetics, as expected for mono and bis-intercalators^9,10^, respectively. YOYO-1 displayed slower dissociation compared to SxO due to an additional, slow, kinetic step (Fig. 1c,d and Supplementary Fig. 1b). The measured kinetics indicate that improvements to BALM can be made by selecting mono-intercalating dyes with high autocatalysis, to optimally match the imaging parameters of the microscope.

**Figure 1.**
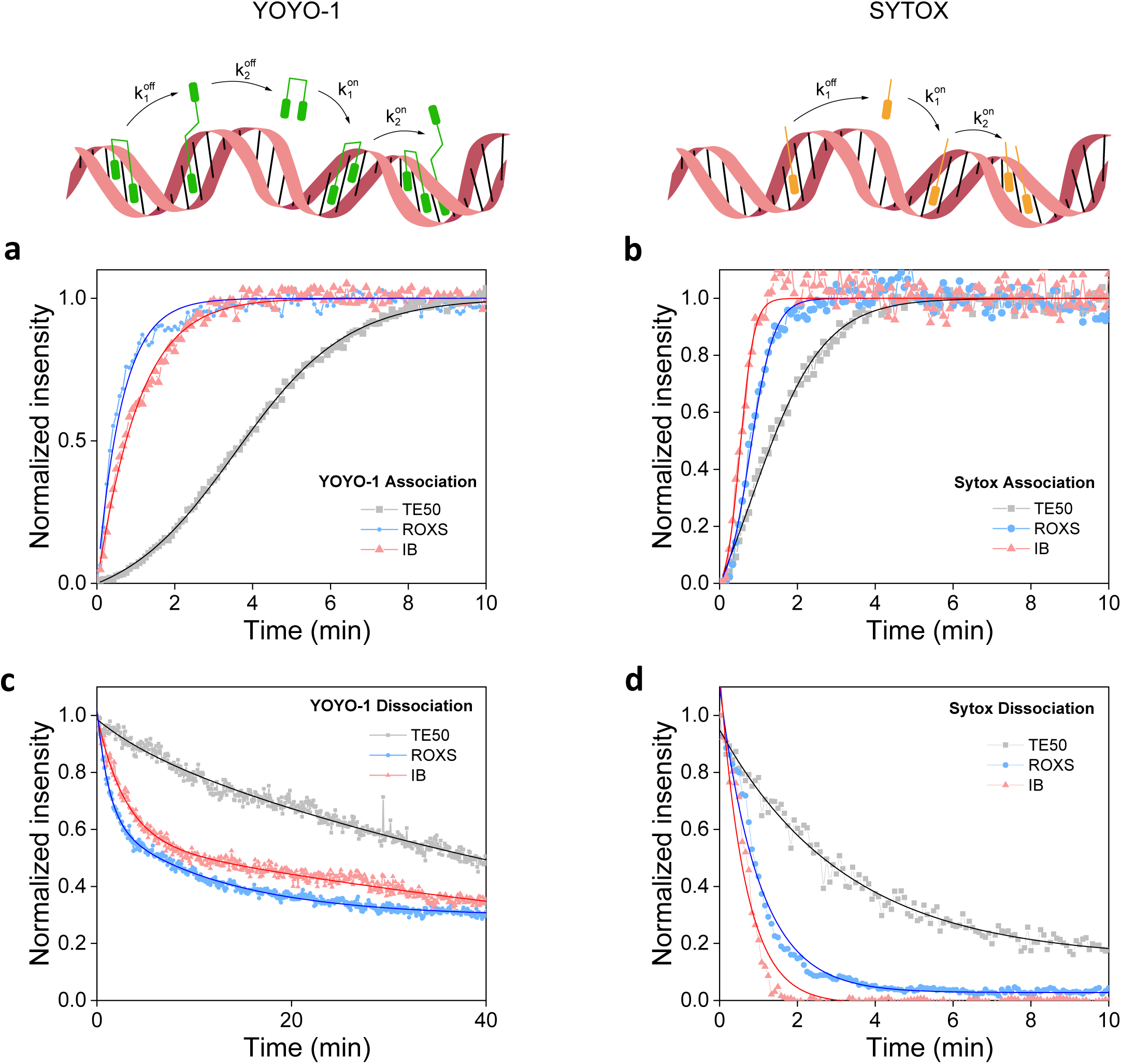
YOYO-1 and SxO binding and unbinding kinetics in different buffers. **a-d**, Time-lapse measurements of association kinetics at 20 nM YOYO-1 (**a**) or SxO (**b**). Dissociation kinetics of YOYO-1 (**c**) and SxO (**d**) under three different buffer conditions. Buffer conditions are TE50 (10 mM Tris pH 8.0, 1 mM EDTA, 50 mM NaCl), TE50 containing either Ascorbic Acid and Methyl Viologen (ROXS) or glucose, glucose oxidase, catalase and MEA-HCl (IB) with concentrations indicated in the methods section. The overlaid lines are fits to the data using the model equations described in the methods section.

Importantly, it was possible to completely remove SxO while more than 30% of YOYO-1 remained on DNA even after extensive washing (Fig. 1c,d) as reported previously^5^. In addition, we found that another oxygen scavenging imaging <u>b</u>uffer (IB; TE50 buffer containing glucose, glucose oxidase, catalase and MEA-HCl)^11^ improved association/dissociation of SxO to a larger extent than ROXS (Fig. 1b,d and Supplementary Fig. 1). These results suggest that SxO in IB should perform best in BALM imaging. To compare the quality of super-resolved images with YOYO-1 in ROXS and SxO in IB, we performed two-dimensional BALM measurements on well-defined DNA origami structures^12^ (Supplementary Fig. 2a-c). Even though it was possible to observe triangular and square-shaped DNA assemblies using both dyes, SxO led to a higher number of localizations than YOYO-1 (Supplementary Fig. 2d).

To investigate 3D DNA architecture using BALM, we introduced optical astigmatism through a cylindrical lens. As the fluorophore point spread function changes depending on the distance from the objective focal plane, astigmatism provides axial position information. This method has been previously applied to <u>st</u>ochastic <u>o</u>ptical reconstruction <u>m</u>icroscopy (STORM)^13^ and DNA PAINT^14,15^. To demonstrate the feasibility of our approach, we first applied 3D BALM to 400 nm diameter microspheres coated with 100 base-pair (bp) long oligonucleotide duplexes (Fig. 2a). DNA-bead conjugates were immobilized onto the surface of a coverslip in a microfluidic flow chamber and 30 pM SxO was introduced in IB. When microspheres were not conjugated to DNA, no binding of SxO to microspheres was observed at this SxO concentration. 3D BALM images of DNA-coated beads showed hollow spherical structures with a mean diameter of 408 ± 14 nm (mean ± std dev) consistent with the expected size of the microspheres (400 nm as measured by the manufacturer) (Fig. 2b,c and Supplementary Video 1).

**Figure 2.**
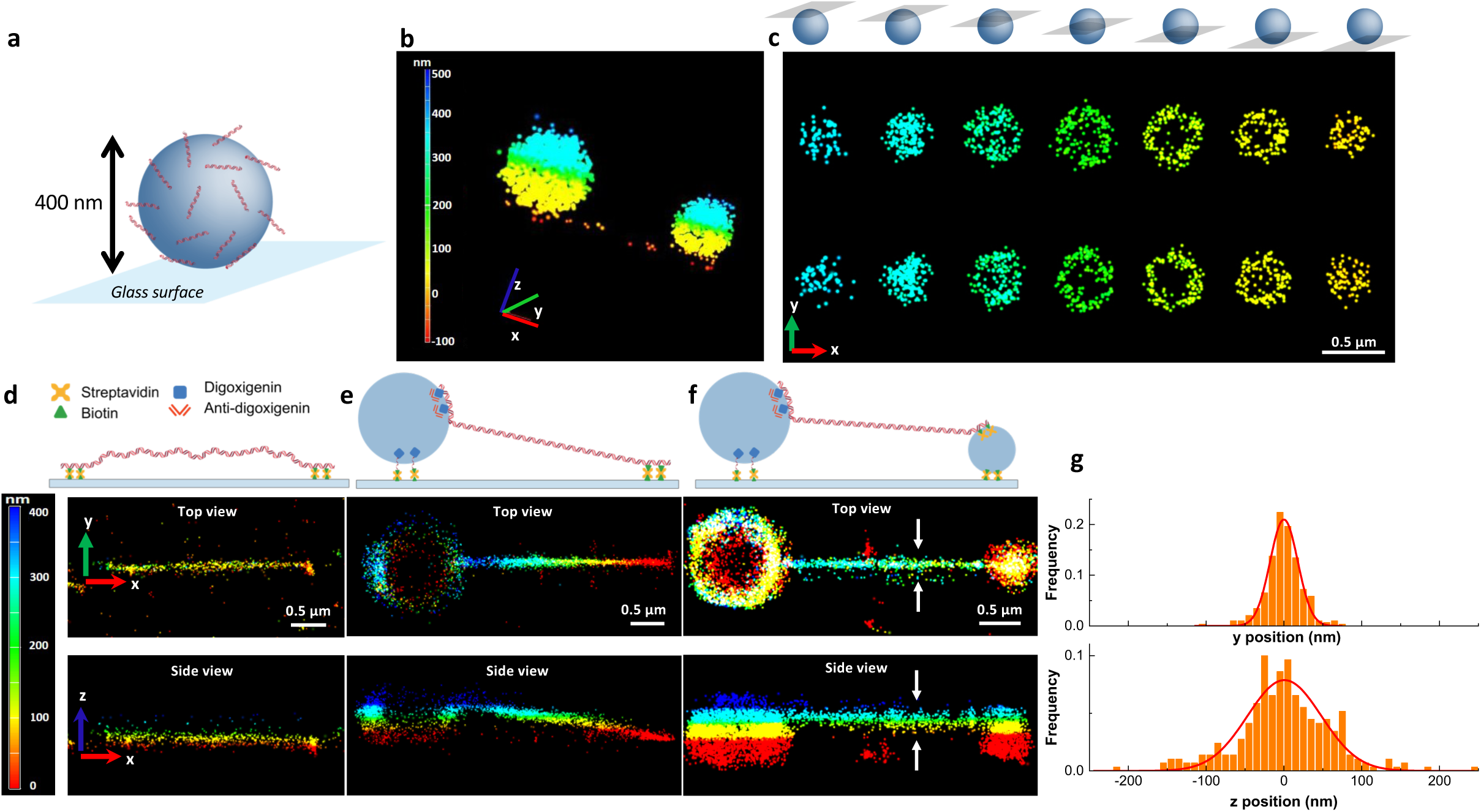
3D BALM imaging of DNA. **a**, A cartoon depiction of a DNA-coated bead immobilized on a glass surface through biotin-streptavidin linkage. **b-c**, 3D BALM images (**b**) and the corresponding cross sections of two DNA-coated microspheres at different axial levels (**c**) with color-coded height map. The schematics above **c** illustrate different z levels. **d-f**, 3D BALM images of 10 kb linear dsDNA tethered to the surface of the glass at both ends (**d**), tethered to the surface at one end and to a 1-µm diameter bead at the other end (**e**), tethered to a 1-µm diameter bead at one end and to a 0.4-µm diameter bead at the other end (**f**). **g**, Line profile for determination of lateral and axial FWHM (representing dsDNA, tethered to a 1-µm diameter bead from one end and to a 0.4-µm diameter bead from the other end). Solids lines are fits to a Gaussian.

To perform 3D BALM on individual DNA molecules, we generated a 10 kilo base pair (kb) linear DNA construct containing biotin modifications on one end and digoxigenin on the opposite end. DNA was tethered to a streptavidin-coated glass coverslip with the biotin-modified end, while the digoxigenin-labelled end of DNA was bound to a 1 µm diameter microsphere conjugated with anti-digoxigenin antibody (Fig. 2e). Microspheres conjugated to DNA were then attached to the surface in the presence of buffer flow to extend the DNA molecules. We performed 3D BALM at relatively higher SxO concentrations (500 pM) due to lower dissociation rates of the dye from highly stretched DNA^8^. At this concentration, SxO aspecifically bound to microspheres making them visible in 3D super-resolution images. Color-coded images of DNA molecules show that the axial position of DNA increases from the surface-tethered end towards the microsphere-tethered end as expected (Fig. 2e, top; Supplementary Fig. 3). An x-z plane projection of a single DNA molecule tethered at both ends to the surface demonstrates that such molecules lie flat on a single z-plane as expected (Fig. 2d). In contrast, a DNA molecule stretched between the surface and a microsphere revealed a clear slope (Fig. 2e, bottom). Due to the hydrodynamic force exerted on microspheres, DNA molecules were stretched to on average 90% of their contour length. A 10 kb DNA molecule (3 µm when stretched to 90% of the contour length) tethered at one end to the surface and at the opposite end to a microsphere 0.5 µm (the radius of the microsphere) above the surface is expected to display a tilt angle of 9.4° from the surface. The average angle of slope was 11 ± 1.7° (n=20, mean ± std dev), consistent with the predicted value. Next, we stretched single DNA molecules between two microspheres thus elevating both ends from the surface. 3D super-resolution images of double-microsphere tethered DNA showed the entire molecule being stretched above the surface (Fig. 2f). The average full width at half maximum of stretched DNA in the lateral, x-y plane was 35.6 ± 1.6 nm (mean ± std dev), higher than previously reported for surface-fixed DNA molecules^5^. Because we tethered DNA to the surface only at their ends, it is likely that thermal fluctuations led to lower resolution compared to completely fixed DNA. Consistently, we found that poorly stretched DNA molecules exhibited larger thickness (Supplementary Fig. 3). The observed axial resolution on DNA stretched between two microspheres was 94.8 ± 5.5 nm (mean ± std dev), consistent with the expected halving of lateral resolution when using the astigmatism in 3D STORM^13^. Together, our results indicate that BALM can be effectively used with optical astigmatism to acquire 3D super-resolution fluorescence images of crowded as well as single DNA molecules.

To explore the applicability of 3D BALM on chromatin, we prepared metaphase chromosome spreads from Jurkat cells. After fixing chromosomes on the glass surface of a flow chamber, we introduced 50 pM SxO in IB and performed BALM imaging. Super-resolution images of SxO-labelled chromosomes yielded an average axial thickness of 714 ± 39 nm (mean ± std dev) (Fig. 3a, Supplementary Video 2, and Supplementary Fig. 4). In addition, structural features of chromosomes such as fine DNA protrusions, that could not be seen with wide-field imaging, were detectable on 3D BALM images.

**Figure 3.**
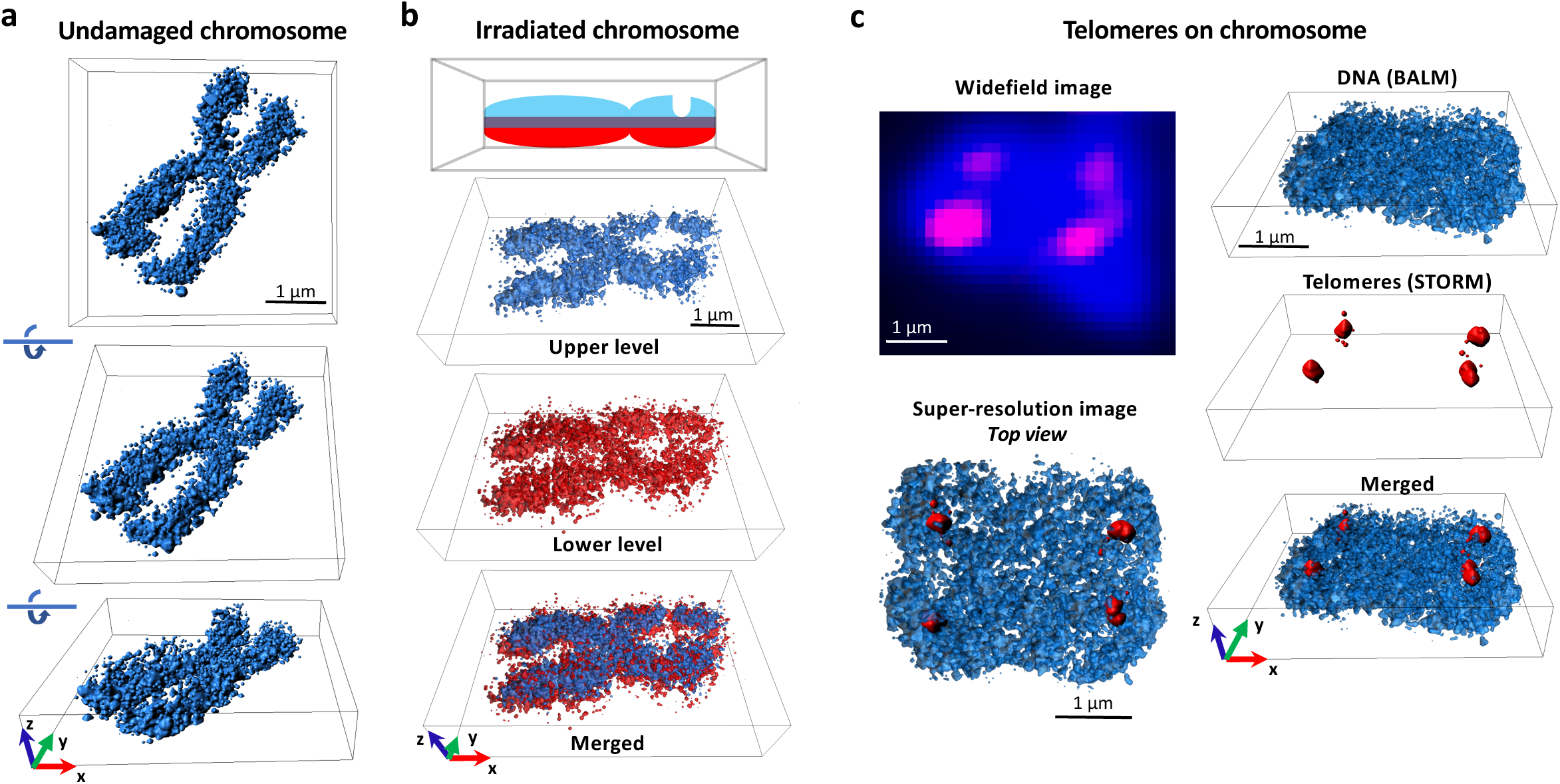
3D super-resolution imaging of metaphase chromosomes. **a**, Three-dimensional BALM images of an undamaged human metaphase chromosome. **b**, 3D BALM images of an irradiated chromosome. A schematic representation of irradiated chromosome with a groove on one of the arms and three-dimensional surface plots at the upper (blue), and lower (red), layers. **c**, (Top left) Wide-field image of a SxO-stained human chromosome (blue) and telomeric regions on the chromosome (magenta). (Bottom left) A top view of merged re-constituted super resolution (BALM) image of the chromosome and re-constituted super resolution (STORM) image of the telomeric regions of this chromosome. (Right) Three-dimensional surface plots of BALM (blue), STORM (red), and both images superimposed.

DNA damaging factors such as ionizing radiation can cause chromosomal abnormalities including chromatid gaps^16^, however the exact nature of these gaps is not clear. To investigate such structures with 3D BALM, we imaged metaphase chromosomes from irradiated cells. Giemsa-stained chromosome spreads displayed small gaps on several chromosomes that would under bright-field microscopy be interpreted as breaks (Supplementary Fig. 5). However, a 3D BALM image of a representative radiation-induced chromatid break (Fig. 3b and Supplementary Video 3) highlights the benefits of 3D BALM for this application as fluorescence density was seen only on lower and not higher z-planes. This suggests that gaps seen in conventional wide-field microscopy may result from thinned chromatin regions rather than complete breaks (further examples in Supplementary Fig. 6). As an alternative approach to introduce chromosome damage, metaphase spreads were treated with sonication in an ultrasonic bath. Similar to irradiated chromosomes, sonication induced thinned regions on some of the chromosomes as revealed by 3D BALM (Supplementary Fig. 7 and Supplementary Video 4).

The ability to differentially target specific DNA sites with fluorescence in situ hybridization (FISH) is a powerful tool to investigate chromosomal arrangements^17^, and has been extensively used in analysis of telomeres^18,19^. We used an Alexa647-labelled protein nucleic acid (PNA) probe to mark telomeric sequences on fixed metaphase chromosomes to obtain 3D organization of PNA-labelled sites with direct STORM^20,21^ (Fig. 3c). Importantly, we were able to acquire 3D BALM images of entire chromosomes and overlay the two super-resolved images (Fig. 3c and Supplementary Video 5). Interestingly, while some telomeric PNA probe localized to the top exterior region on chromosomes (Fig. 3c), others bound to the interior regions and adopted extended shapes (Supplementary Fig. 8). With this method, it will be possible to analyse telomeric structures in detail under different stress conditions.

We have demonstrated the use of fluorescent DNA intercalators for 3D super-resolution imaging of isolated DNA molecules as well as chromatin. Thus, 3D BALM is a versatile method for investigating the architecture of chromosomes. In the future, other DNA binding fluorescent dyes can be tested for specific applications such as cell-permeable fluorophores for 3D BALM in living cells^22^. We provide a kinetic analysis to allow careful selection of intercalators, the concentration, and the buffer conditions necessary. We demonstrated the ability to combine 3D BALM and STORM making it possible to superimpose three-dimensional super-resolution images of DNA with other chromosome components. Therefore, super-resolution images of other factors such as γH2AX marking DNA double-strand breaks^23^ can be acquired and overlaid onto 3D BALM images.

## Acknowledgements

We thank Raffaella Carzaniga and Lucy Collinson from the electron microscopy science technology platform at the Francis Crick Institute for acquiring TEM images of DNA origami structures. This work was supported by the Francis Crick Institute, which receives its core funding from Cancer Research UK (FC001221), the UK Medical Research Council (FC001221), and the Wellcome Trust (FC001221). D.R.B. received funding from the European Union’s Horizon 2020 research and innovation programme under the Marie Skłodowska-Curie grant agreement No 657479. S.Y.A.T. was supported by the Academy of Medical Sciences (SBF001\1019) and the Wellcome/EPSRC Centre for Medical Engineering at King’s College London (WT 203148/Z/16/Z).

## Author contributions

S.Y., D.R.B., S.Y.A.T, and H.Y. conceived and designed the project. S.Y.A.T. prepared metaphase chromosome samples. S.Y. performed all other experiments. S.Y., D.R.B., and H.Y. interpreted the data and wrote the paper with input from S.Y.A.T.

## Methods

### Flow cell preparation

Flow cells were prepared essentially as described previously^24^. Briefly, glass coverslips (24×40 mm, 0.17 ±0.01 mm, VWR, 630-2746) were cleaned with two rounds of sonication in ethanol and 1M potassium hydroxide 30 minutes each round, treated with (3-Aminopropyl)triethoxysilane in acetone and subsequently functionalized with a mixture of polyethylene glycol (PEG)-Succinimidyl Valerate (SVA) (MW 5000, Laysan Bio Inc.) and Biotin-PEG-SVA (MW 5000, Laysan Bio Inc.) in 100 mM sodium bicarbonate buffer (pH 8.2). A flow channel was prepared by sandwiching double-sided tape between the functionalized coverslip and a glass slide (VWR, 48300-025) containing two holes. An inlet (Intramedic, PE20; inner diameter 0.015”, outer diameter 0.048”) and an outlet (Intramedic, PE20; inner diameter 0.015”, outer diameter 0.048”) tubing are attached to the holes on the glass slide and sealed with epoxy (Devcon, 5 minute Epoxy). The inlet tubing was submerged into an Eppendorf tube filled with buffer, while the outlet tubing was attached to an automated syringe pump (Harvard Apparatus, Pump 11 Plus Single Syringe) to withdraw buffer through the flow channel.

### Preparation of DNA constructs

#### One-end biotinylated λ DNA for measurement of dye kinetics

To measure association and dissociation kinetics of SxO and YOYO-1, one end of linear λ phage DNA (New England Biolabs) was labelled with biotin. To this end, a ∼0.5 kb PCR substrate was generated from pUC19 vector with primers 5’-ATGCCGGGAGCAGACAAGCCCGTC-3’ and 5’Phosphate-AGGTCGCCGCCCGGAAG-AGCAGCTGGCACGACAGGTTTCCCG-3’ in the presence of Biotin-16-dUTP (Enzo). Biotin-modified PCR substrate was then nicked with Nt.BspQI (New England Biolabs) generating a 12-nt 3’ tail complementary to one end of λ DNA. 10-fold excess of biotinylated PCR was mixed with λ DNA, ligated with T4 DNA ligase and further purified by gel electrophoresis. To functionalize the surface with streptavidin, 0.2 mg/ml streptavidin in phosphate buffered saline (PBS) was introduced into the flow channel, incubated for 5 min, and free streptavidin was removed by washing the flow cell with blocking buffer (20 mM Tris-HCl pH 7.5, 50 mM NaCl, 2 mM EDTA, 0.2 mg/ml BSA). 1 nM biotin-modified λ was introduced in blocking buffer, incubated for 5-10 min, and excess DNA was removed with blocking buffer. To stretch and fix DNA on the surface, Label IT® Biotin (Mirus) in water was flown at 100 µl/min for 5 minutes. While one end of Label IT® covalently crosslinks to λ DNA, the opposite biotin end binds to the streptavidinated surface, thus immobilizing the DNA on the surface.

#### DNA origami

Nanoscale DNA origami was built as described in ref^12,25^. A single-stranded scaffold DNA, and over 200 oligonucleotides, called staple strands, were used for self-assembly of DNA molecules into triangular and rectangular DNA origami shape with the edge length of 120 nm. A mixture was prepared with 100 nM staple mix and 10 nM scaffold from M13 bacteriophage (Tilibit) in folding buffer (5 mM Tris pH 8.0, 1 mM EDTA, 5 mM NaCl, 20 mM MgCl_2_). This mixture was annealed in a PCR thermocycler using a fast-linear cooling step from 80°C to 65°C over 1 hour, followed by 42 hours linear cooling ramp from 65°C to 24°C. Annealed sample was then subjected to gel electrophoresis in 0.5x TBE buffer in the presence of 10 mM MgCl_2_ at 75 V for 3 hours in a cold room. Finally, DNA origami was gel extracted via electroelution and concentrated using spin columns (Amicon Ultra 0.5 ml, 3kDa MWCO). To immobilize DNA origami on the surface, coverslips were streptavidin functionalized as described above. Label IT® Biotin was introduced into the flow cell and incubated for 10 min. After washing the flow cell with folding buffer, DNA origami was introduced in folding buffer, incubated for a period of time (2-30 min) to obtain sufficient surface coverage, and excess origami was removed with folding buffer. All buffers used in downstream applications contained 10 mM MgCl_2_ to prevent disassembly of origami structures.

#### DNA-coated microspheres

A 100-bp duplex DNA containing a biotin at one end was prepared by mixing equimolar amounts of oligonucleotides 5’GGTTGGCGAATTCCCATTGCCTCAGCATCCGGTACCTCAGCACGACGTTGTAAAACGAGCCTTCACCGTGGTGAGTTTGTCTTCTCGAAGCAGTCAACCA3’BiotinTEG and 5’TGGTTGACTGCTTCGAGAAGACAAACTCACCACGGTGAAGGCTCGTTTTACAACGTCGTGCTGAGGTACCGGATGCTGAGGCAATGGGAATTCGCCAACC3’ (Integrated DNA Technologies Inc.) in buffer (10 mM Tris pH 8.0, 100 mM NaCl and 1 mM EDTA), heating to 85°C and slowly cooling down to room temperature. Streptavidin-coated polystyrene particles (0.4 µm diameter, Spherotech Inc.) were washed twice in T50 buffer (10 mM Tris pH 8.0, 1mM EDTA, 50 mM NaCl) by centrifugation and sonicated in an ultrasonic water bath for one minute to break down aggregates. DNA and microspheres were mixed in T50 buffer, incubated overnight at room temperature on a rotator, and unbound DNA was removed by washing the microspheres twice with T50.

#### Biotin/digoxigenin-modified 10 kb linear DNA

λ DNA was digested with ApaI (New England Biolabs), separated on a 0.5% agarose gel, and the resulting 10 kb fragment was isolated via gel extraction (Qiagen). Biotinylation of one end of this fragment was achieved using the same protocol described for biotin-modification of λ DNA. To label the other end of 10kb DNA with digoxigenin, a ∼0.5 kb fragment from pUC19 was PCR amplified with 5’ATGCCGGGAGCAGACAAGCCCGTC3’ and 5’ATGGGCCCAGCTGGCACGACAGG-TTTCCCG3’ in the presence of digoxigenin-11-dUTP (Roche). The digoxigenin-labeled DNA was digested with ApaI, gel purified, mixed in 10-fold excess with biotin-modified 10 kb DNA, and ligated using T4 DNA ligase (NEB). The substrate was separated on a 0.5% agarose gel and purified by gel extraction.

10 kb DNA was immobilized on the surface from its biotinylated end as described above. To attach the other end of DNA to microspheres, we conjugated polyclonal anti-digoxigenin antibody (anti-dig, Roche) to carboxylated polystyrene particles (1 µm diameter, Spherotech Inc.). Anti-dig-coated microspheres in blocking buffer were introduced into the flow cell and allowed at least 1 hr to bind DNA. Free microspheres were removed by washing the chamber with blocking buffer. To stretch DNA and attach microspheres to the surface, an oligonucleotide (10 nM in blocking buffer) containing digoxigenin on one side and biotin on the other (5’BiotinTEG-TTTTTTTTTTTTTTTTTTTTTTTTTTTTTT3’Digoxigenin) was withdrawn at 250 µl/min for one minute.

To stretch DNA between two microspheres, 0.4 µm-diameter streptavidin coated beads were attached on the PEG-Biotin-modified coverslip surface of the flow cell. DNA was then introduced to attach the streptavidin-coated microspheres from the biotinylated end of DNA. Because surface is not coated with streptavidin at this stage, DNA binds only to streptavidin-conjugated microspheres and not to the surface. After removing free DNA, 1 µm-diameter anti-dig-conjugated microspheres were introduced for binding to the free digoxigenin-end of DNA molecules. After removing free anti-dig-coated microspheres, streptavidin was introduced to coat the PEG-biotin surface. Finally, free streptavidin was removed, and biotin/digoxigenin double-labelled oligonucleotide was introduced as described above to attach anti-dig-conjugated microspheres to streptavidin-functionalized surface.

#### Metaphase chromosome spreads

Human T lymphocytic Jurkat cells (ATCC) were grown in suspension at 37°C in a humidified atmosphere with 5% CO_2_ in RPMI-1640 medium (Sigma-Aldrich) supplemented with 1.5 mM L-glutamine (PAA Laboratories, Austria), 10% fetal bovine serum (Thermo Fisher) and penicillin/streptomycin (50units/ml, 50µg/ml) (Thermo Fisher). Cells were maintained between 1 and 20 × 10^5^ cells/ml. Cells in medium (1.5 × 10^6^ cells/mL; 5.5 mL in 15 mL centrifuge tube) were irradiated at 1 Gy or sham-irradiated at room temperature by a ^137^Cs source at 5 Gy/min. Samples were kept on ice to and from the irradiator. Cells were then incubated with colcemid (0.15 mg/mL; Sigma-Aldrich) at 37°C in a humidified atmosphere with 5% CO_2_ for 1.5 hours. Medium was subsequently replaced with 2.5 mL 75 mM KCl hypotonic buffer after spinning cells at 500 x g for 5 minutes at room temperature. Ten minutes later, cells were spun as before and fixed in 3:1 methanol:acetic acid for 10 minutes at room temperature. After a final spin, cells were resuspended in 3:1 methanol:acetic acid at 22.5 × 10^6^ cells/mL (375 mL) and kept at −20°C until used for studies. 10 µl of chromosome sample was drop from 40 cm height onto a glass coverslip that was sonicated only in ethanol. Glass slide was kept at room temperature for 15 minutes to dry and a flow chamber was assembled such that chromosomes would stay inside the flow channel. To induce mechanical DNA damage, the glass coverslip with metaphase spread was treated with 20 minutes sonication in an ultrasonic water bath before assembly of the flow chamber.

To stain telomeric regions, coverslip containing dried chromosome sample was placed into 4% paraformaldehyde in PBS for 10 minutes and rinsed with PBS. The sample was first treated with RNase A (0.1 mg/ml in PBS) at 37°C for 15 minutes, then with pepsin (0.002% in 100 mM HCl) at 37°C for 15 minutes. Between and after RNase A and pepsin treatments the sample was washed with PBS. Next, the coverslip was washed in series with 70%, 85%, and 100% ethanol for two minutes each. Finally, the sample was incubated with 200 nM of TelC-Cy5 (Panagene) in probe solution (20 mM Tris-HCl pH 7.5, 10% goat serum (Thermo Fisher)) overnight at room temperature, in the dark, for hybridization. The sample was submersed in buffer (100 mM sodium phosphate dibasic, 50 mM sodium phosphate monobasic dehydrate, 0.1% Triton X-100) at 37°C for 5 minutes and washed with PBS at room temperature. A flow chamber was then prepared with the coverslip containing PNA-labeled chromosomes.

### Imaging and Data analysis

Samples were imaged using Nikon Ti-E microscope with 100x oil NA 1.49 SR Apo TIRF objective and Andor iXon EMCCD camera. Images were captured with NIS-Elements AR software equipped with N-STORM module. YOYO-1 or SxO was imaged with 488- or 532-nm lasers, respectively. For dye association/dissociation measurements, images were acquired with 5 s time-lapse, 150 ms exposure time, 300 EM gain at 1% of the maximum laser intensity. When imaging dissociation kinetics, DNA was first stained with 20 nM YOYO-1 or SxO in TE50 buffer (10 mM Tris pH 8.0, 1 mM EDTA, 50 mM NaCl) for 30min. After binding and unbinding data acquisition, 150 frames without any time delay between frames were acquired for photobleaching controls. All data were taken at room temperature.

In BALM measurements, a field of view comprising 256 × 256 pixels was acquired with the EMCCD camera. BALM imaging was performed with maximum 532 nm laser intensity at 50-500 pM SxO in IB (TE50 (10 mM Tris pH 8.0, 1mM EDTA, 50 mM NaCl) supplemented with 40 mg/ml glucose, 0.05 mg/ml glucose oxidase (Sigma-Aldrich), 1 µg/ml catalase (Sigma-Aldrich) and 100 mM MEA-HCl (Sigma-Aldrich)) or with the maximum 488 nm laser intensity at 200 pM YOYO-1 in T50 buffer containing another oxygen scavenging system ROXS consisting of 10 mM Ascorbic Acid (Sigma-Aldrich) and 1mM Methyl Viologen hydrate (Acros Organics). STORM imaging was performed with the maximum 647 nm laser intensity in IB. When performing BALM on DNA origami samples, origami buffer (50 mM Tris, 50 mM NaCl, 1 mM EDTA, 10 mM MgCl_2_) was used instead of T50 buffer. For drift correction, 0.19 µm diameter Flash Red microspheres (Bangs Laboratories) were used as fiducial markers for BALM with SxO, and 0.1 µm diameter yellow/green microspheres (FluoSpheres, Thermo Fisher) were used for BALM with YOYO-1 and for STORM.

10,000-60,000 frames were acquired without delay with 50 ms exposure for BALM and 10 ms exposure for STORM imaging. For 3D BALM/STORM, 3D-Stack function of N-STORM module was used to acquire images at multiple different Z-positions over the time with 200 nm step size. A set of sequentially acquired images was analysed by Nikon N-STORM software (NIS-Elements AR) and formed into a super-resolution image. Super-resolution images include information such as position (in the X and Y axis direction is referred to 2D-STORM/BALM image, in the x, y and z axis direction is referred to 3D-STORM/BALM image), size, and intensity of each individual fluorescent molecule. Imaris (Bitplane) was used to generate surface plots from 3D BALM/STORM super-resolution reconstituted images.

#### Association/Dissociation kinetics

The dissociation as a function of time data was modelled as a multi-step process with intermediate states existing between the bound and free dye states. Formally, this is described by a sum of exponential decays^26,27^, with the number chosen through minimising chi-squared. YOYO-1 dissociation is best described by the reaction

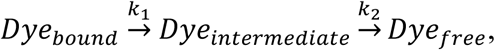

and so, the intensity by a double exponential decay

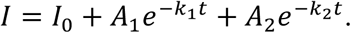

SxO is best described with no intermediate states

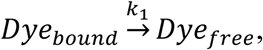

and so, by a single exponential decay

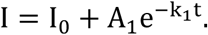

The association of both dyes follows a logistic curve as a function of time and we model the data as an autocatalytic reaction

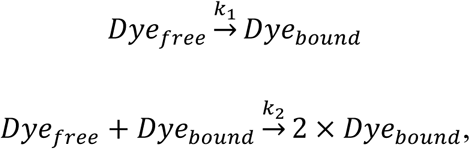

and so the intensity by^28^

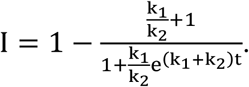

## Supplementary Video Legends

**Supplementary Video 1. 3D BALM video of DNA-conjugated 400-nm diameter microspheres.** This video corresponds to the images shown in Figure 2b-c.

**Supplementary Video 2. 3D BALM video of an undamaged chromosome represented as surface plot.** This video corresponds to the chromosome shown in Figure 3a.

**Supplementary Video 3. 3D BALM video of an irradiated chromosome represented as surface plot.** This video corresponds to the chromosome shown in Figure 3b. The images were collected at two different z-levels: high (blue), low (red).

**Supplementary Video 4. 3D BALM video of a sonication-treated chromosome represented as surface plot.** This video corresponds to the chromosome shown in Supplementary Figure 7.

**Supplementary Video 5. 3D super-resolution video of a chromosome and telomeric regions.** This video corresponds to the chromosome shown in Figure 3c. 3D BALM (blue, DNA) and STORM (red, telomeric PNA) images merged.

## References

1 Martin, M., Terradas, M., Hernandez, L. & Genesca, A. gammaH2AX foci on apparently intact mitotic chromosomes: not signatures of misrejoining events but signals of unresolved DNA damage. Cell Cycle 13, 3026–3036, (2014).

2 Ushiki, T. & Hoshi, O. Atomic force microscopy for imaging human metaphase chromosomes. Chromosome Res 16, 383–396, (2008).

3 Daban, J. R. Electron microscopy and atomic force microscopy studies of chromatin and metaphase chromosome structure. Micron 42, 733–750, (2011).

4 Patterson, G., Davidson, M., Manley, S. & Lippincott-Schwartz, J. Superresolution imaging using single-molecule localization. Annu Rev Phys Chem 61, 345–367, (2010).

5 Schoen, I., Ries, J., Klotzsch, E., Ewers, H. & Vogel, V. Binding-activated localization microscopy of DNA structures. Nano Lett 11, 4008–4011, (2011).

6 Yardimci, H., Loveland, A. B., Habuchi, S., van Oijen, A. M. & Walter, J. C. Uncoupling of sister replisomes during eukaryotic DNA replication. Mol Cell 40, 834–840, (2010).

7 Yardimci, H. et al. Bypass of a protein barrier by a replicative DNA helicase. Nature 492, 205–209, (2012).

8 Biebricher, A. S. et al. The impact of DNA intercalators on DNA and DNA-processing enzymes elucidated through force-dependent binding kinetics. Nat Commun 6, 7304, (2015).

9 Pyle, J. R. & Chen, J. Photobleaching of YOYO-1 in super-resolution single DNA fluorescence imaging. Beilstein journal of nanotechnology 8, 2296–2306, (2017).

10 Almaqwashi, A. A., Paramanathan, T., Rouzina, I. & Williams, M. C. Mechanisms of small molecule–DNA interactions probed by single-molecule force spectroscopy. Nucleic Acids Research 44, 3971–3988, (2016).

11 Metcalf, D. J., Edwards, R., Kumarswami, N. & Knight, A. E. Test samples for optimizing STORM super-resolution microscopy. J Vis Exp, (2013).

12 Dietz, H., Douglas, S. M. & Shih, W. M. Folding DNA into twisted and curved nanoscale shapes. Science 325, 725–730, (2009).

13 Huang, B., Wang, W., Bates, M. & Zhuang, X. Three-dimensional super-resolution imaging by stochastic optical reconstruction microscopy. Science 319, 810–813, (2008).

14 Iinuma, R. et al. Polyhedra self assembled from DNA tripods and characterized with 3D DNA-PAINT. Science 344, 65–69, (2014).

15 Jungmann, R. et al. Multiplexed 3D cellular super-resolution imaging with DNA-PAINT and Exchange-PAINT. Nat Methods 11, 313–318, (2014).

16 Durkin, S. G. & Glover, T. W. Chromosome fragile sites. Annu Rev Genet 41, 169–192, (2007).

17 Levsky, J. M. & Singer, R. H. Fluorescence in situ hybridization: past, present and future. J Cell Sci 116, 2833–2838, (2003).

18 Baerlocher, G. M., Vulto, I., de Jong, G. & Lansdorp, P. M. Flow cytometry and FISH to measure the average length of telomeres (flow FISH). Nat Protoc 1, 2365–2376, (2006).

19 Sfeir, A. & de Lange, T. Removal of shelterin reveals the telomere end-protection problem. Science 336, 593–597, (2012).

20 Wombacher, R. et al. Live-cell super-resolution imaging with trimethoprim conjugates. Nat Methods 7, 717–719, (2010).

21 Genet, M. D., Cartwright, I. M. & Kato, T. A. Direct DNA and PNA probe binding to telomeric regions without classical in situ hybridization. Molecular Cytogenetics 6, 42, (2013).

22 Zhang, X. et al. A targetable fluorescent probe for dSTORM super-resolution imaging of live cell nucleus DNA. Chemical Communications 55, 1951–1954, (2019).

23 Valdiglesias, V., Giunta, S., Fenech, M., Neri, M. & Bonassi, S. gammaH2AX as a marker of DNA double strand breaks and genomic instability in human population studies. Mutat Res 753, 24–40, (2013).

24 Tanner, N. A. & van Oijen, A. M. in Single Molecule Tools, Part B:Super-Resolution, Particle Tracking, Multiparameter, and Force Based Methods Methods in Enzymology 259–278 (2010).

25 Rothemund, P. W. Folding DNA to create nanoscale shapes and patterns. Nature 440, 297–302, (2006).

26 House, J. E. Principles of chemical kinetics. (Wm. C. Brown, 1997).

27 Tiwari, P. B., Wang, X., He, J. & Darici, Y. Analyzing surface plasmon resonance data: choosing a correct biphasic model for interpretation. Rev Sci Instrum 86, 035001, (2015).

28 Watzky, M. A. & Finke, R. G. Transition Metal Nanocluster Formation Kinetic and Mechanistic Studies. A New Mechanism When Hydrogen Is the Reductant: Slow, Continuous Nucleation and Fast Autocatalytic Surface Growth. Journal of the American Chemical Society 119, 10382–10400, (1997).

