## Supplementary Figures for "Three-dimensional super-resolution fluorescence imaging of DNA"

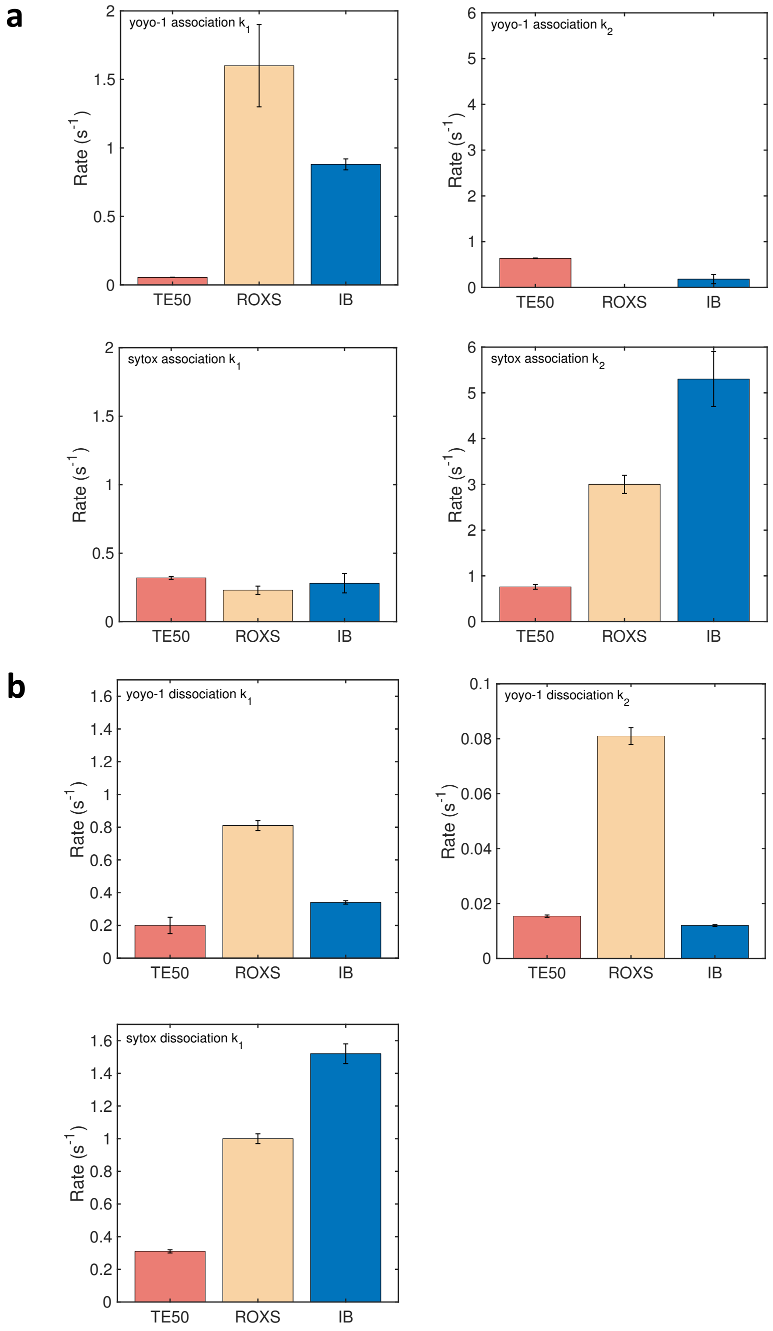


**Supplementary Figure 1. Comparison of YOYO-1 and SxO association and dissociation kinetics. a,** Kinetic parameters of association. Despite YOYO-1 having a higher rate of reaction for free dye to DNA bound dye, $k_1$, the much faster autocatalysis of DNA bound dye, $k_2$, results in faster association for SxO. **b,** Kinetic parameters of dissociation. Dissociation of SxO is faster mostly due to the lack of a second, slow, kinetic step.

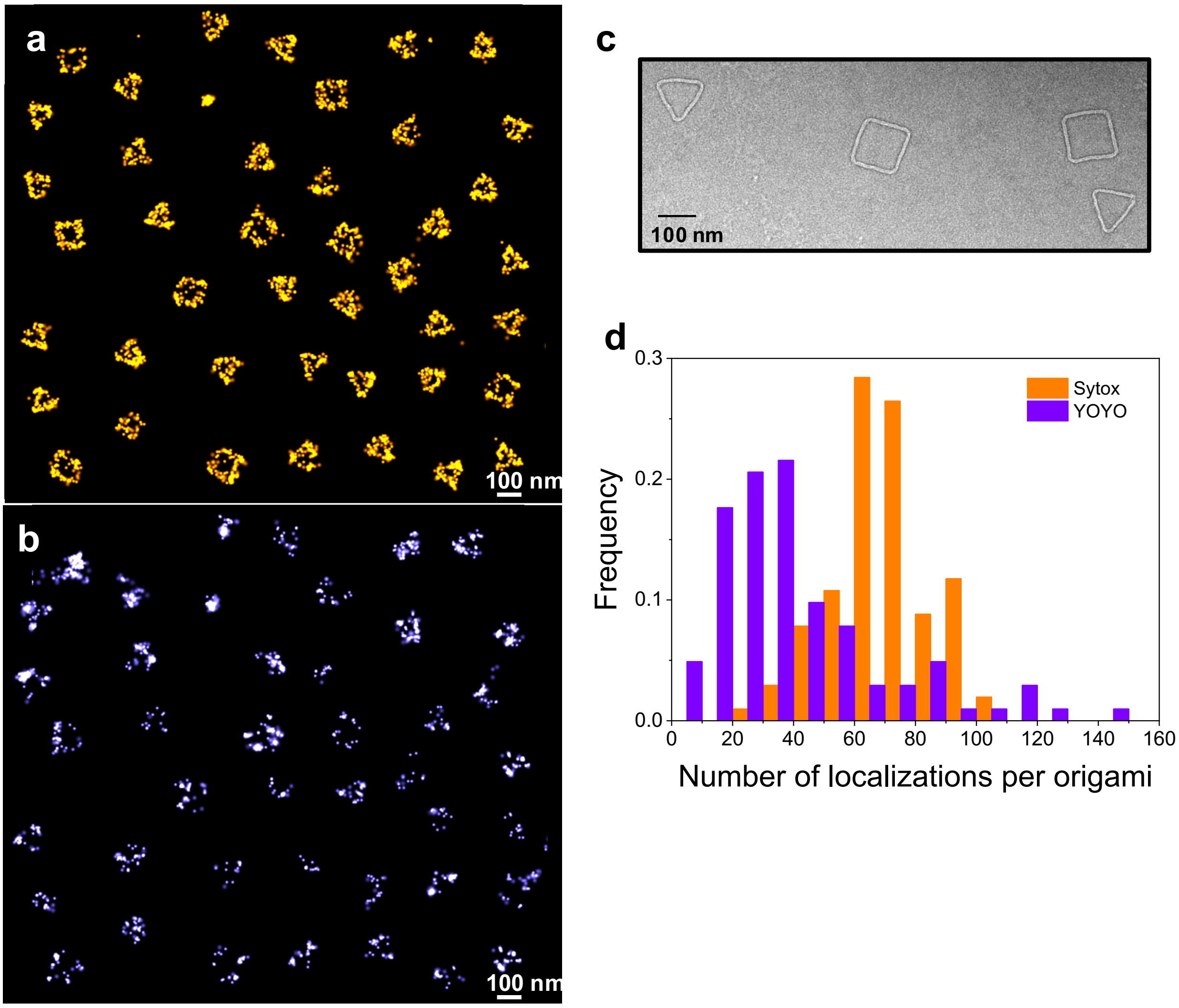


**Supplementary Figure 2. 2D BALM images of DNA origami with edge length of 120 nm. a-b,** Super-resolution imaging of triangular and square-shaped DNA origami with DNA-intercalating dye SxO (**a**) or YOYO-1 (**b**). The images were constructed using molecules from multiple fields of view. **c,** Transmission electron microscopy images of DNA origami molecules. **d,** Distributions of number of localizations per DNA origami molecule, frequency, reconstituted with 50 pM SxO or 200 pM YOYO-1.

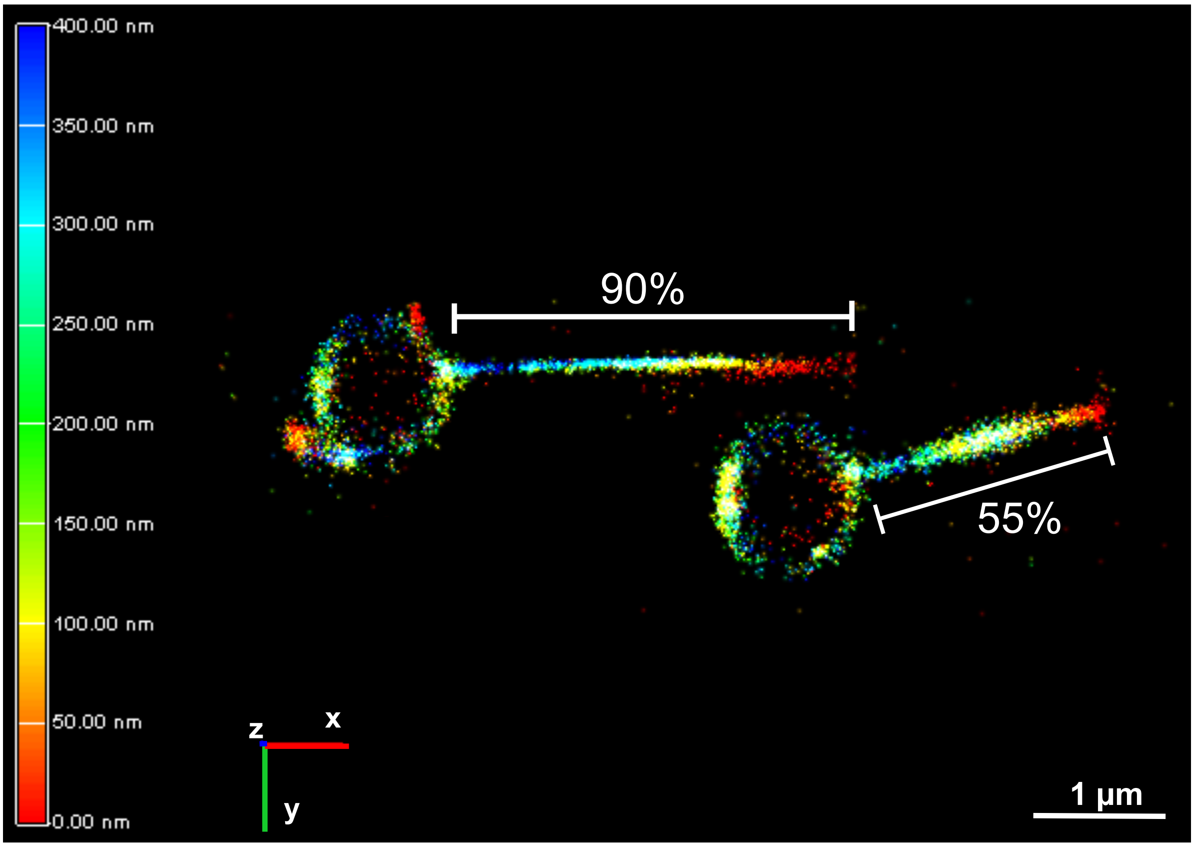


**Supplementary Figure 3. 3D BALM images of linear DNA molecules.** BALM images of 10 kb DNA constructs stretched between the surface of the glass slide and 1 μm bead. The width of a DNA molecule stretched to 55% of its contour length is larger than that of 90% stretched DNA due to thermal fluctuations of the DNA molecule.

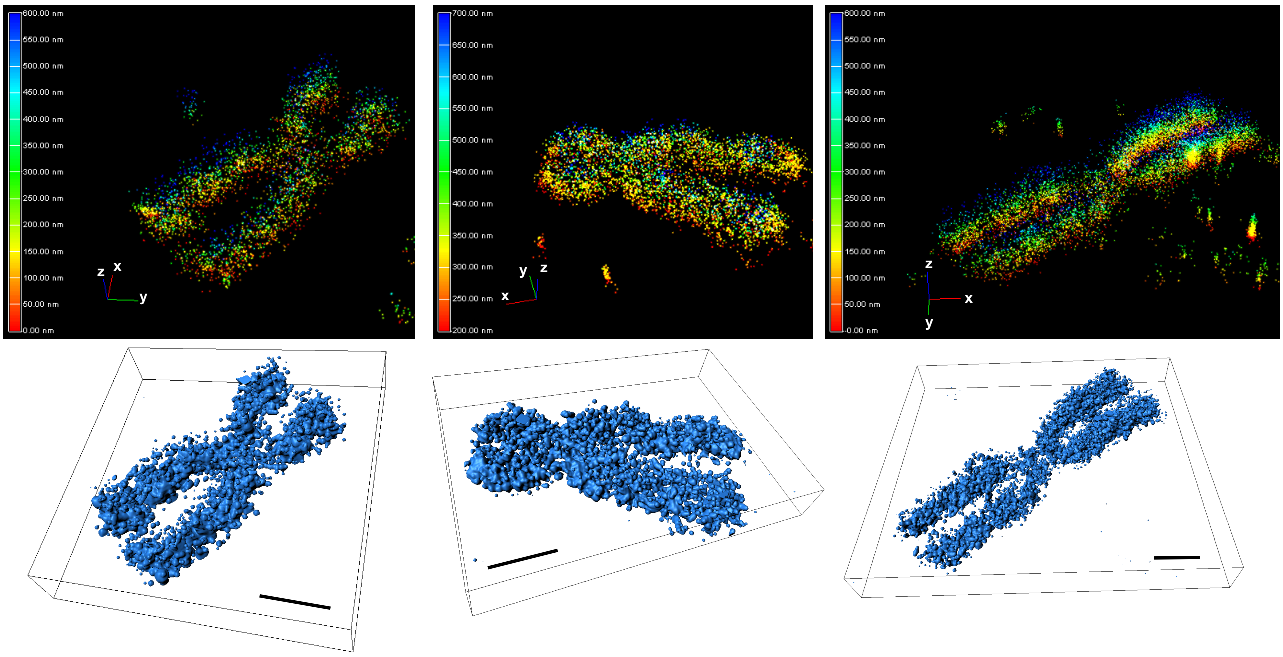


**Supplementary Figure 4. 3D BALM images of metaphase chromosomes.** Three-dimensional BALM images of human metaphase chromosomes and (bottom) representing three-dimensional surface plots. Scale bars: 1 µm.

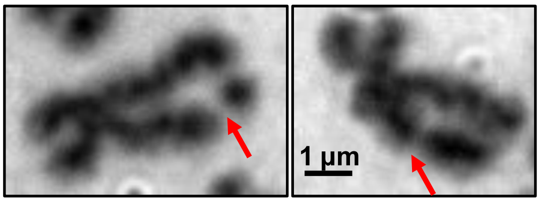


**Supplementary Figure 5. Sample metaphase chromosomes with chromatid breaks.** Brightfield images of 1Gy irradiated metaphase chromosomes. Red arrows show the chromatid gaps.

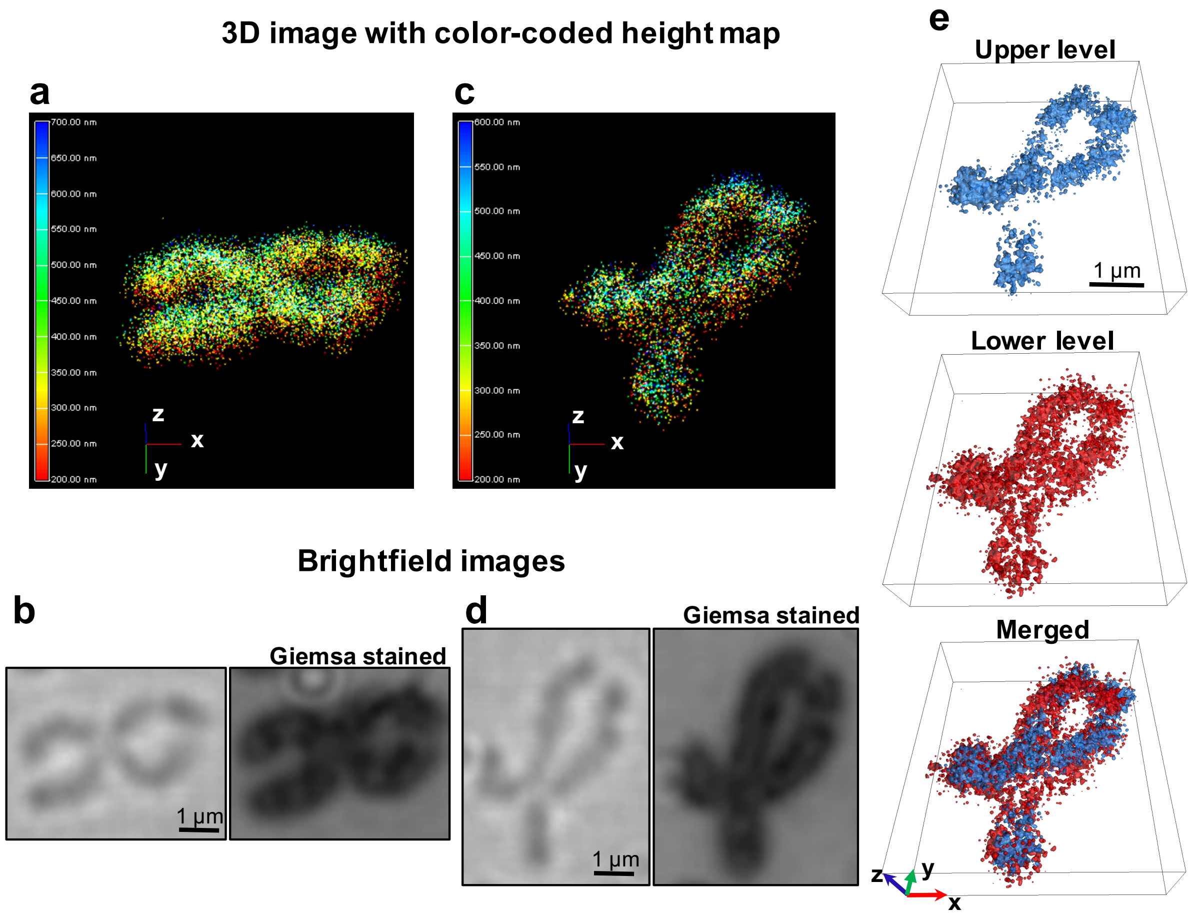


**Supplementary Figure 6. 3D super-resolution imaging of irradiated metaphase chromosomes.** **a**, Three-dimensional BALM image of human chromosome (same chromosome shown in Fig. 2b) with a break as a result of radiation with color-coded height map and **b**, the corresponding brightfield image of the same chromosome before (left) and after (right) Giemsa staining. **c,** Another example 3D BALM image of an irradiated human chromosome containing a gap and **d,** its brightfield image of before (left) and after (right) Giemsa staining. **e,** Representing three-dimensional surface plot of the chromosome in **c** at upper (blue), lower (red) z planes and merged images.

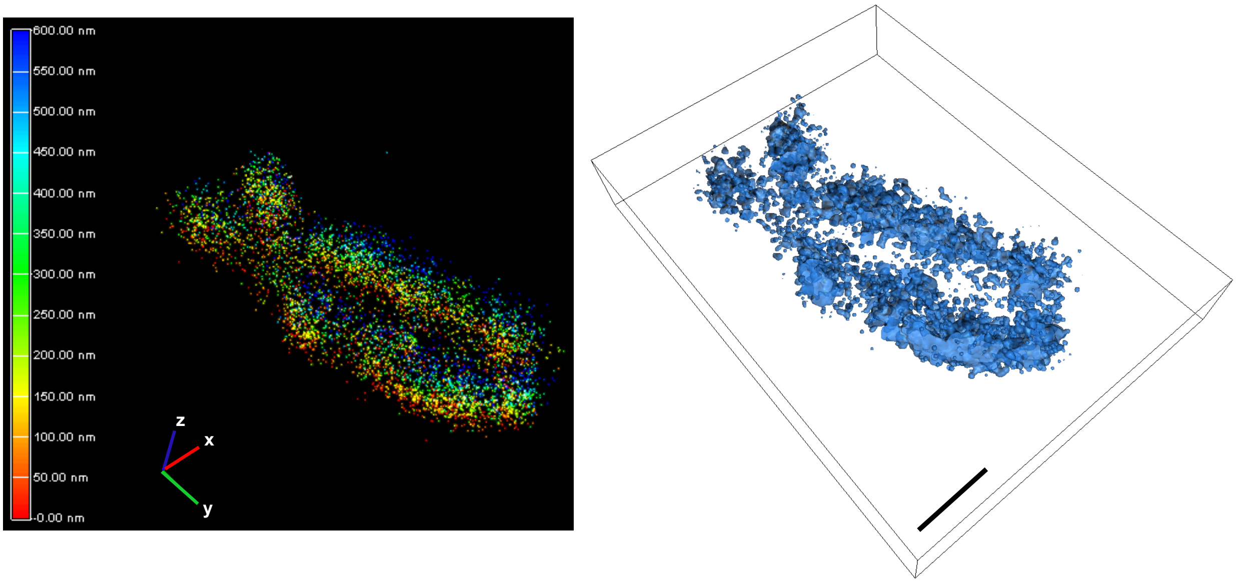


**Supplementary Figure 7: 3D BALM imaging of sonication-induced DNA damage.** 3D BALM image of a sonication-treated metaphase chromosome (left) and the corresponding three-dimensional surface plot (right). Scale bar: 1 µm.

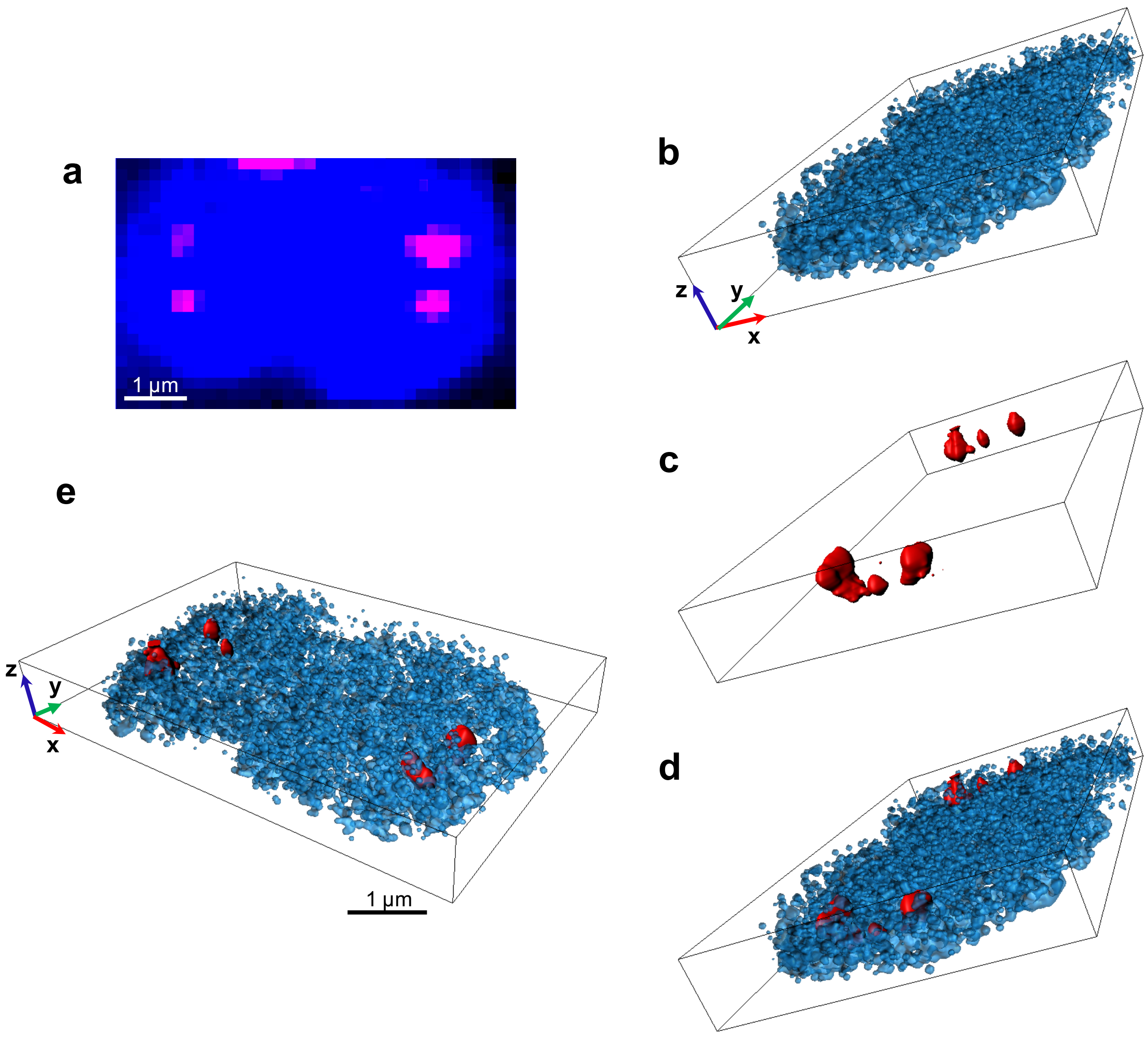


**Supplementary Figure 8. 3D BALM on metaphase human chromosomes and 3D STORM on telomeric regions. a,** Merged wide-field image of a human metaphase chromosome (blue) and telomeric regions (magenta). **b,** 3D BALM image of the same chromosome. **c,** 3D STORM image of telomeric regions on the chromosome. **d-e,** Composite 3D BALM and STORM images of the chromosome and the telomeric regions on the chromosome.

| | $I_0$ | $k_1$ | $k_2$ | $A_1$ | $A_2$ |
|---|---|---|---|---|---|
| YOYO Dissociation | | | | | |
| TE50 | - | 0.20 ± 0.05 | 0.0154 ± 0.0004 | 0.071 ± 0.010 | 0.92 ± 0.01 |
| ROXS | 0.294 ± 0.002 | 0.81 ± 0.03 | 0.081 ± 0.003 | 0.387 ± 0.007 | 0.341 ± 0.005 |
| IB | - | 0.34 ± 0.01 | 0.0120 ± 0.0003 | 0.439 ± 0.006 | 0.561 ± 0.004 |
| YOYO Binding | | | | | |
| TE50 | | 0.055 ± 0.001 | 0.636 ± 0.008 | | |
| ROXS | | 1.6 ± 0.3 | 6E-7 ± 0.7 | | |
| IB | | 0.88 ± 0.04 | 0.18 ± 0.10 | | |
| SYTOX Dissociation | | | | | |
| TE50 | 0.15 ± 0.010 | 0.31 ± 0.01 | | 0.80 ± 0.01 | |
| ROXS | 0.028 ± 0.004 | 1.00 ± 0.03 | | 1.07 ± 0.02 | |
| IB | -0.011 ± 0.005 | 1.52 ± 0.06 | | 1.16 ± 0.03 | |
| SYTOX Binding | | | | | |
| TE50 | | 0.32 ± 0.01 | 0.76 ± 0.05 | | |
| ROXS | | 0.23 ± 0.03 | 3.0 ± 0.2 | | |
| IB | | 0.28 ± 0.07 | 5.3 ± 0.6 | | |

**Supplementary Table 1.** Values of kinetic parameters from the model fits in Figure 1.
